# Protein synthesis rates and ribosome occupancies reveal determinants of translation elongation rates

**DOI:** 10.1101/465914

**Authors:** Andrea Riba, Noemi Di Nanni, Nitish Mittal, Erik Arhné, Alexander Schmidt, Mihaela Zavolan

**Affiliations:** Institut de Génétique et de Biologie Moléculaire et Cellulaire, Université de Strasbourg, 1 rue Laurent Fries 67404, Illkirch CEDEX, France; Institute of Biomedical Technologies, National Research Council, Segrate (MI), Italy; Department of Industrial and Information Engineering, University of Pavia, Italy; Biozentrum, University of Basel, Klingelbergstrasse 70, 4056 Basel, Switzerland

## Abstract

Although protein synthesis dynamics has been studied both with theoretical models and by profiling ribosome footprints, the determinants of ribosome flux along open reading frames (ORFs) are not fully understood. Combining measurements of protein synthesis rate with ribosome footprinting data, we here inferred translation initiation and elongation rates for over a thousand ORFs in exponentially-growing wildtype yeast cells. We found that the amino acid composition of synthesized proteins is as important a determinant of translation elongation rate as parameters related to codon and tRNA adaptation. We did not find evidence of ribosome collisions curbing the protein output of yeast transcripts, either in high translation conditions associated with exponential growth, or in strains in which deletion of individual ribosomal protein genes leads to globally increased or decreased translation. Slow translation elongation is characteristic of RP-encoding transcripts, which have markedly lower protein output than other transcripts with equally high ribosome densities.

**Significance Statement:** Although sequencing of ribosome footprints has uncovered new aspects of mRNA translation, the determinants of ribosome flux remain incompletely understood. Combining ribosome footprint data with measurements of protein synthesis rates, we here inferred translation initiation and elongation rates for over a thousand ORFs in yeast strains with varying translation capacity. We found that the translation elongation rate varies up to ~20-fold among yeast transcripts, and is significantly correlated with the rate of translation initiation. Furthermore, the amino acid composition of synthesized proteins impacts the rate of translation elongation to the same extent as measures of codon and tRNA adaptation. Transcripts encoding ribosomal proteins are translated especially slow, having markedly lower protein output than other transcripts with equally high ribosome densities.

## Introduction

Gene expression analysis frequently relies on the high-throughput sequencing of cellular mRNAs. While the mRNA expression levels may be sufficient to decipher how cells respond to specific stimuli, they explain protein abundances only to a limited extent, with coefficients of determination R^2^ in the range of 0.14-0.41 (1, 2). Protein levels vary over a much wider range than the levels of the corresponding mRNAs, indicating extensive regulation of protein metabolism, and especially synthesis (1). Translation is predominantly regulated at the initiation step (3), whose rate varies broadly between mRNAs, depending on the structural accessibility of the 5’ end to translation factors, and on the presence of upstream open reading frames (uORFs). The latter generally hinder translation of the main ORF (4). Translation elongation rates also differ between mRNAs, primarily due to differences in tRNA availability. Whether and how the translation elongation rate is dynamically modulated is currently debated (2, 5–8). TRNA availability, translational co-folding of the polypeptide chain and the presence of positively charged amino acids in the nascent peptide have all been linked to variation in elongation rate (5–8). Furthermore, it has been proposed that the codon usage is the substrate of ‘translational programs’ that adjust the protein output of of specific classes of mRNAs to the state (proliferation or differentiation) of the cell (9). However, explicit comparison of the coverage of 5’ and 3’ halves of ORFs by ribosome footprints did not reveal clear differences, indicating that bottlenecks in elongation along coding regions are uncommon (2).

Insights into the dynamics of translation and putative bottlenecks have emerged from theoretical studies, in particular of the totally asymmetric simple exclusion process (TASEP), introduced almost five decades ago (10). In a simple form of this model, ribosomes bind to mRNAs according to an initiation rate, move stochastically to downstream codons with an average elongation rate, if these codons are not already occupied by ribosomes, and are released at the end of the coding region with a given termination rate. The interplay of these rates gives rise to three distinct regimes. If termination is at least as fast as elongation, ribosome collisions within ORFs never become relevant. However, when termination is slow relative to elongation and initiation is fast, ribosome start to ‘collide’ and the protein output drops markedly. Otherwise, as long as elongation is not limiting, the rate of protein synthesis increases linearly with the rate of translation initiation, in parallel with the density of ribosomes on the open reading frame (ORF) (11).

These predictions can be tested with currently available technologies, as the TASEP model makes specific predictions about the relationship between ribosome flux (protein synthesis rate) and ribosome density along ORFs. Ribosome density along ORFs can be studied with high resolution by sequencing of ribosome-protected mRNA footprints, a method known as ribosome profiling (2). The approach has already uncovered novel principles of resource allocation and translation regulation (12, 13). Furthermore, model-based analyses of ribosome profiling data uncovered sources of local variation in ribosome densities and translation elongation along transcripts (14). However, the ribosome flux has rarely been measured directly, in spite of mass spectrometry-based methods being able to provide estimates of synthesis and degradation rates for a substantial fraction of eukaryotic proteomes (15). Direct measurement of protein synthesis rate is necessary to detect global changes in translation capacity between conditions (16) and for studying translation in an ORF-specific manner, as ribosome profiles can be used to infer protein synthesis rate only up to a constant factor.

To fill this gap and further uncover factors that underlie variations in translation elongation rates between ORFs we measured protein synthesis rates transcriptome-wide, by pulsed stable isotope labeling of amino acids in culture (pSILAC), in the widely-studied experimental model of exponentially growing yeast cells. Combined analysis of pSILAC and ribosome footprinting data revealed the range of variation in translation elongation rates between yeast ORFs. Among broadly studied determinants of this rate, most indicative were the availability of cognate tRNAs and the frequency of positively charged amino acids in the synthesized protein. We found no evidence that translation is curbed by ribosome collisions either in exponentially growing wildtype yeast or in mutant strains with global alterations in translation. Rather, we found that mRNAs encoding positively charged proteins, in particular ribosomal proteins, are translated at a markedly slower rate than other mRNAs with similar ribosome densities.

## Results

### Ribosome allocation is largely explained by the copy number and length of open reading frames

To uncover determinants of translation speed in exponentially growing yeast cells, we analyzed a recently published ribosome footprinting data set obtained in this system (4), from the perspective of the TASEP model of translation (Figure 1A). Denoting by the *N* number of ribosomes bound to a coding region of *L* codons, and assuming that the rate with which a ribosome completes the polypeptide chain is given by the product of the frequency of finding a ribosome at the stop codon 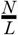,and the effective ribosome translocation (and termination) rate *k_el_*, the change in the number of ribosomes bound on the mRNA is given by the differntial equation 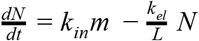. Here *m* is the number of mRNA molecules, *k_in_* is the effective rate of translation initiation on an mRNA molecule, and we assume the broad region of parameter values where ribosome collisions are rare. This model predicts that the number of ribosome-protected fragments (RPFs) mapping to a specific mRNA is proportional to the mRNA abundance, the length of the ORF and the ratio of the effective rates of initiation and elongation, 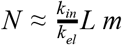.

**Figure 1.**
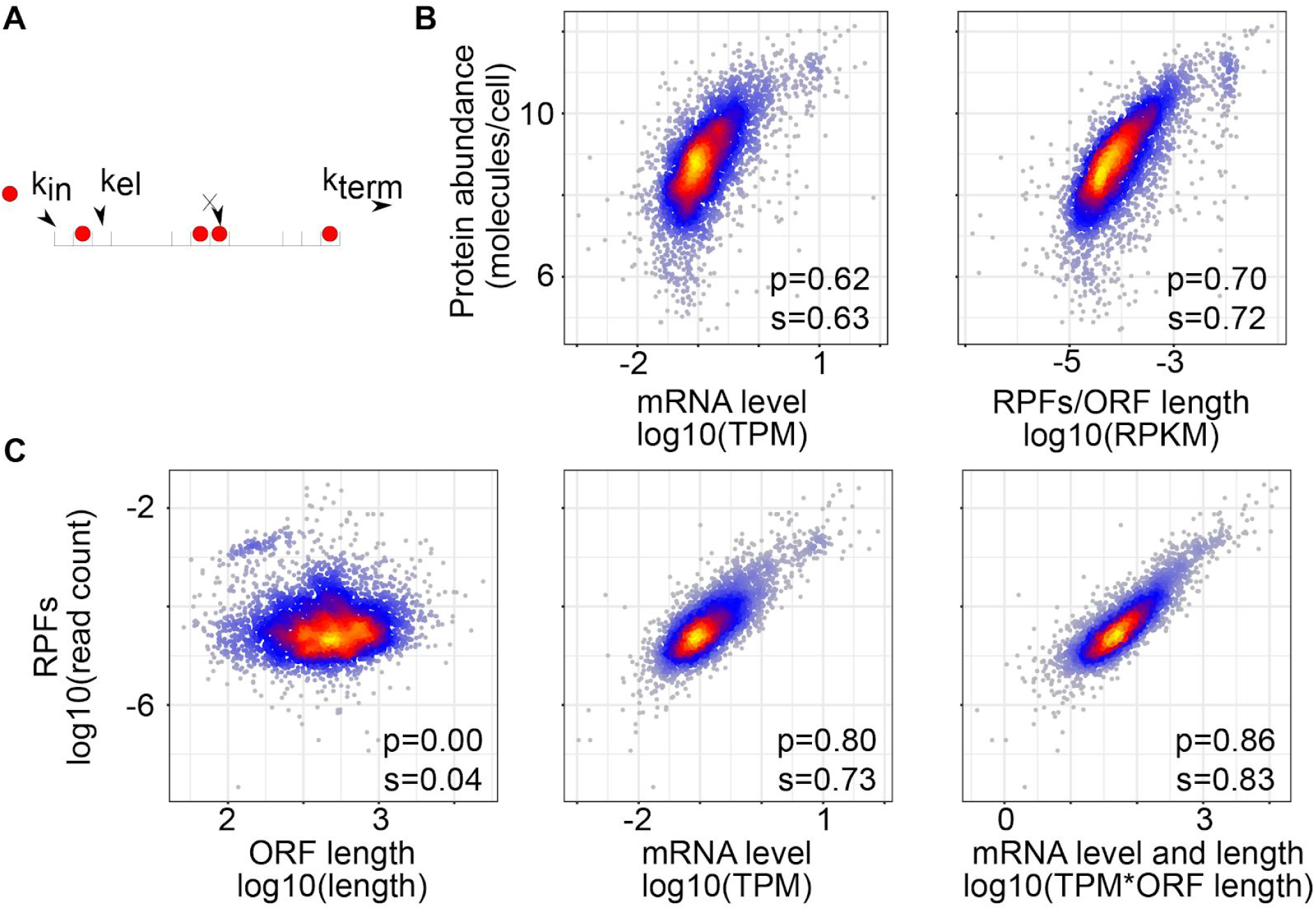
Predicted and observed relationships in gene expression in the *BY4741* yeast strain. **(A)** Illustration of the classical Totally Asymmetric Exclusion Process (TASEP) with constant rates of initiation, elongation and termination. **(B)** Relationship between protein abundance (18) and the density of RPFs on the ORF or the mRNA abundance. **(C)** Relationship between the number of RPFs mapped to individual mRNAs and the corresponding ORF length, mRNA level, and both.

Testing this prediction with the previously published experimental data set (4), we found that the number of RPFs mapped to a specific mRNA indeed correlated very well with the relative abundance of the mRNA estimated by mRNA sequencing. However, further incorporating the ORF length slightly reduced rather than improved the correlation (Supplementary Figure 1). As it was reported that the estimation of mRNA abundance by RNA sequencing is in fact a critical parameter to control in the analysis of ribosome footprinting data (4), we repeated the analysis of scaling behavior with estimates of mRNA abundance obtained by sequencing of RNAs purified directly with oligo(dT) from yeast cell lysates (17). Of note, the 3’-end bias in ORF coverage by RNA-seq reads, that strongly affects the accuracy of mRNA abundance estimates (4), was quite similar between the two data sets (Supplementary Figure 2). We found that when using the RNA samples obtained by oligo(dT)-based purification, both the mRNA level and ORF length contributed to the number of RPFs, as expected (Figure 1B). We therefore used this mRNA sequencing data set for the analyses described below, but present similar results with the RNA-seq data from (4) in the Supplementary material.

The mRNA levels alone explained 65% of the variance in RPF numbers. Further taking into account the ORF length increased this number to 74% (Figure 1B), setting an upper bound of 25% on the variance in RPFs that could be due to differences in ribosome density along transcripts. From previously published measurements of protein levels in the same yeast strain (18), we further inferred that the number of RPFs explained ~50% of the variance in protein levels, compared to only 36% explained by mRNA abundance (Figure 1C).

### Ribosome allocation predicts protein synthesis rates

Theoretical analysis of the TASEP model showed that the main dynamical regimes are defined by the density and the flux of ribosomes on transcripts (11), the latter corresponding to the protein synthesis rate. To infer the translation regime of individual yeast mRNAs, we determined relative ribosome densities on individual ORFs from the RPF and RNA-seq data, knowing the ORF lengths. The estimates that we obtained here correlated quite well (Figure 2A, Spearman correlation coefficient = 0.46, p-value = 1.6e-194) with those from a much earlier study that determined the distribution of individual mRNA species across polysomal fractions corresponding to one, two, three etc. translating ribosomes, with microarrays (19). Having computed ribosome densities for each ORF, we used pulsed stable isotope labeling by amino acids in cell culture (pSILAC) to measure the corresponding protein synthesis rates.

**Figure 2.**
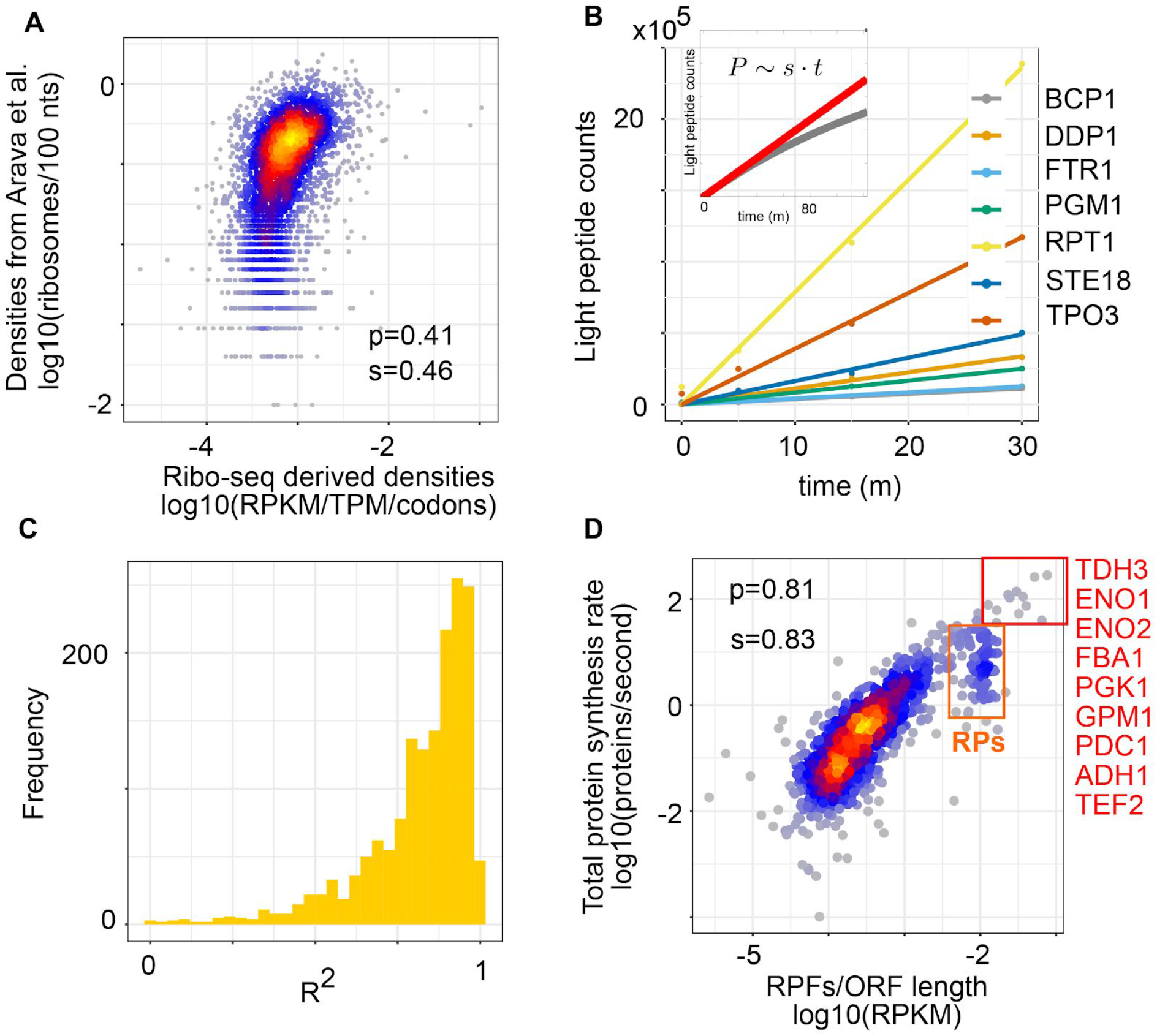
Analysis of protein synthesis rates. **(A)** Ribosome densities derived from the sequencing of RPFs (x-axis) or estimated based on the relative abundance of RNAs across polysomal fractions by (19) (y-axis). **(B)** Protein synthesis rates *s* can be estimated from the dynamics of light peptide (*P*) accumulation within a short time interval *t* after medium change (inset). Example of linear fits to the peptide accumulation curves for the proteins indicated in the legend. **(C)** Histogram of R^2^ values of the linear fit for all 1’616 measured proteins. **(D)** Relationship between ribosome allocation per codon and the protein synthesis rate. Highlighted in the red box are the proteins with highest synthesis rates. The orange box highlights the cluster of RPs.

On a short time scale, upon shifting cells from a medium with heavy isotope-containing amino acids to a medium with light isotope-containing peptides, ‘light’ peptides should accumulate proportionally to the protein synthesis rates (Figure 2B). Indeed, we found that the light peptide accumulation in the first 30 minutes after the medium change was very well described by a linear model (R^2^ for the linear fit > 0.8 for 1’114 of the 1’616 proteins, Figure 2C). Furthermore, the protein synthesis rates thus estimated correlated better with the number of RPFs than with the protein levels (Pearson correlation coefficients of 0.81 and 0.7, respectively, Figure 2D and Figure 1C). This conforms to the expectation that RPFs reflect protein synthesis, while protein levels are set by the balance between synthesis and degradation.

### Protein synthesis rates are not limited by ribosome collisions in exponentially growing yeast cells

Although protein synthesis rates increased linearly with the ribosome allocation over the entire range, a small cluster of ORFs did not conform to this relation, but had distinctly lower protein output than other ORFs with similarly high ribosome densities (Figure 2D, orange box). These ORFs encoded almost exclusively ribosomal proteins, while other highly translated ORFs, with a single exception (the translation elongation factor 2, TEF2), encoded genes involved in sugar metabolism (Figure 2D, red box). Indeed, Gene Ontology terms Glucose metabolic process and Gluconeogenesis and KEGG Glycolysis/Gluconeogenesis pathway were strongly enriched in this set (false discovery rate for these biological processes: 1e-14 and 2e-12, respectively, Fig. 2D). To understand the dynamics of translation on individual ORFs, we then sought to infer their absolute rates of translation initiation and elongation.

A yeast cell needs about 2 hours to divide (20) and contains about 5 x 10^7^ protein molecules (21), whose average half lives are ~8.8 hours (22). These numbers define the total number of proteins produced by a yeast cell per unit time, allowing us to convert the relative protein synthesis rates inferred from the pSILAC time series to absolute rates of molecules per unit time. Further taking into account the estimated number of 40’000 (23, 24) mRNA molecules in a yeast cell, we obtained protein synthesis rates per mRNA (see Methods). To directly compare the experimental data (Figure 3A) with predictions of the TASEP model (Figure 3B), we converted the relative ribosome densities that we obtained from sequencing of RPFs to absolute densities of ribosomes per codon (rpc) using the first principal component of the scatter of RPF-based densities as a function of absolute density measured by (19) (Figure 2A). Protein synthesis rate - ribosome density results for individual ORFs, computed using either oligo(dT) or Ribo-zero RNA sequencing data sets, are given in Supplementary Tables 1 and 2.

**Figure 3.**
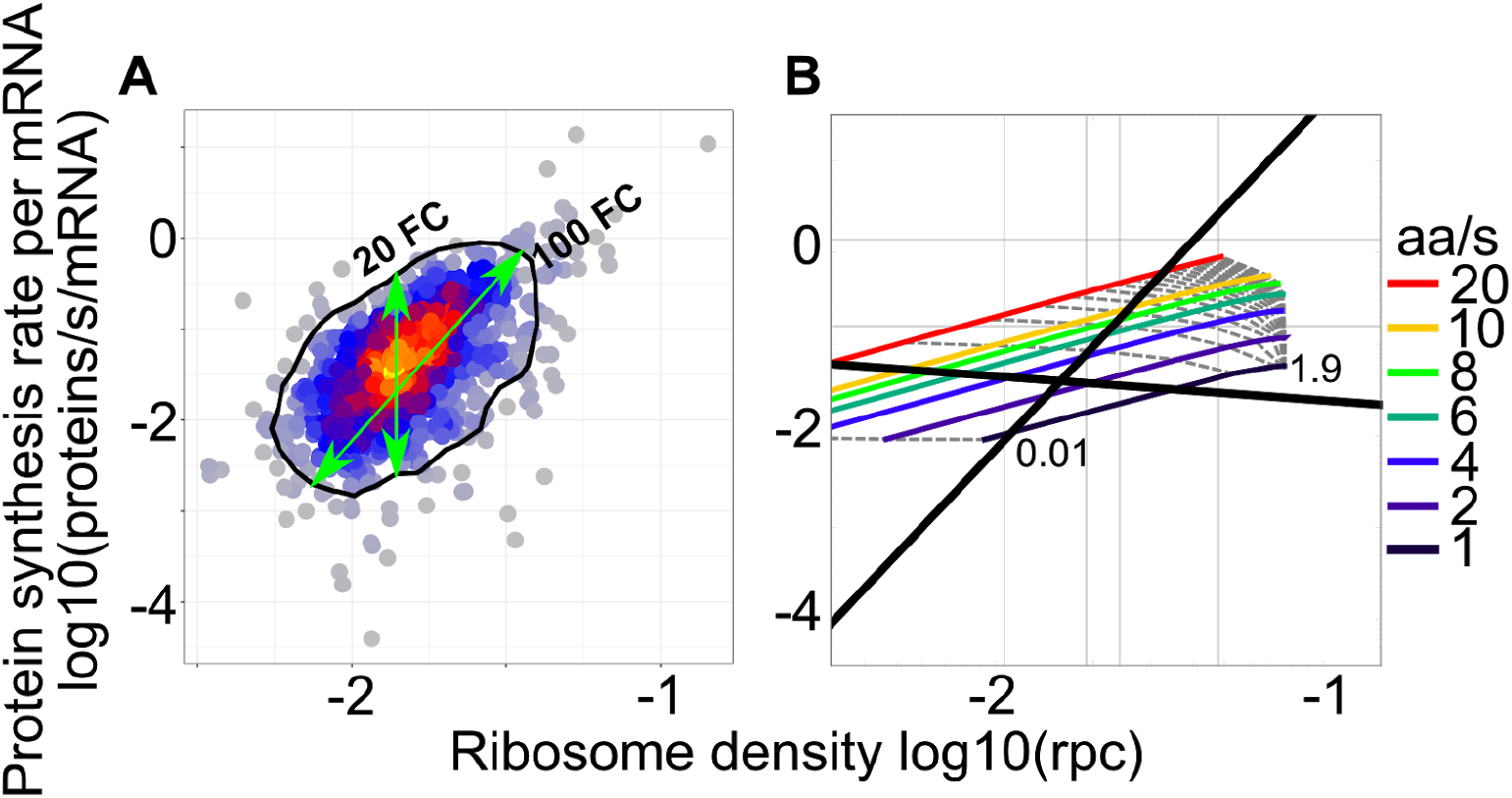
Observed and predicted relationship between the protein synthesis rate and ribosome density on the corresponding ORF. **(A)** Experimental results: protein synthesis rates were measured by pSILAC and converted to molecules per mRNA per second from the expected protein mass doubling time; ribosomes densities were obtained from the fit of ribosome footprint densities to numbers of ribosomes per codons (rpc) estimated by (19). The black contour indicates 90% of the empirical distribution approximated through the two-dimensional Kernel Density Estimation from the R package ‘MASS’. **(B)** TASEP model predictions with isoclines corresponding to individual translation initiation (gray dotted lines, rate range: 0.01 - 1.9/*s*, increments of 0.1 starting from second line at 0.1, mean 0.04/*s*) and elongation rates (colored lines, rate range: 1 - 20 aa/s, mean 2.63 aa/*s*). Superimposed are the two principal components of the experimental data shown in (A).

In the model, ORFs that initiate translation at very low rates have very low protein output and their ribosome coverage per codon reflects the rate of translation elongation (Figure 3B). Although our data set contained only few proteins with very low synthesis rate, the ~10-fold range that we infer this way is comparable to the 20-fold range that we observe for ORFs with the same RPF density (Figure 3A). The model also predicts that protein synthesis rate and ribosome coverage increase linearly with the initiation rate, as long as ribosome collisions do not halt elongation (Figure 3B). The ~100-fold range of variation in protein output in the experimental data corresponds to a similar range of variation in translation initiation rate. Thus, our analysis indicates that translation is primarily regulated at the level of initiation, as reported before (4). The regime of high ribosome density and low protein output exhibited by the model was not observed in our data (Figure 3A), indicating that ribosome collisions do not curb the rate of protein synthesis in the high protein output regime of exponentially growing yeast.

### Combining protein synthesis rate and ribosome footprint density reveals the speed of translation elongation

If ribosome queueing events are rare, proteins should be synthesized at the rate at which new chains are initiated *s* ≃ *k _in_*, the ribosome density can be approximated as 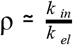, and the ratio of protein synthesis rate to ribosome density (SDR) will effectively be the translation elongation rate, 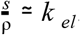. We used this relationship to uncover features of the mRNA or of the corresponding protein that most strongly affect elongation (Figure 4). The positive correlation of the SDR with the average speed of ribosomes, calculated from the normalized ribosome densities at individual codons, served as control (Figure 4A). We found that all measures related to tRNA availability and codon usage (tRNA adaptation index (tAI) (25), ‘normalized tAI’ (26), fraction of optimal codons (FOP) (27) and codon adaptation index (CAI) (28)) (see Methods for details) correlated positively with the SDR, as expected (29) (Figure 4A). Interestingly, the ribosome density was also positively correlated with the SDR (Figure 4A). This could be due to ribosomes promoting translation speed by resolving RNA secondary structure (see also below). Participation of the newly incorporated amino acid in a protein domain was also associated with faster elongation, and this association was not due to a specific type of domain such as *α*-helix or *β*-sheet.

**Figure 4.**
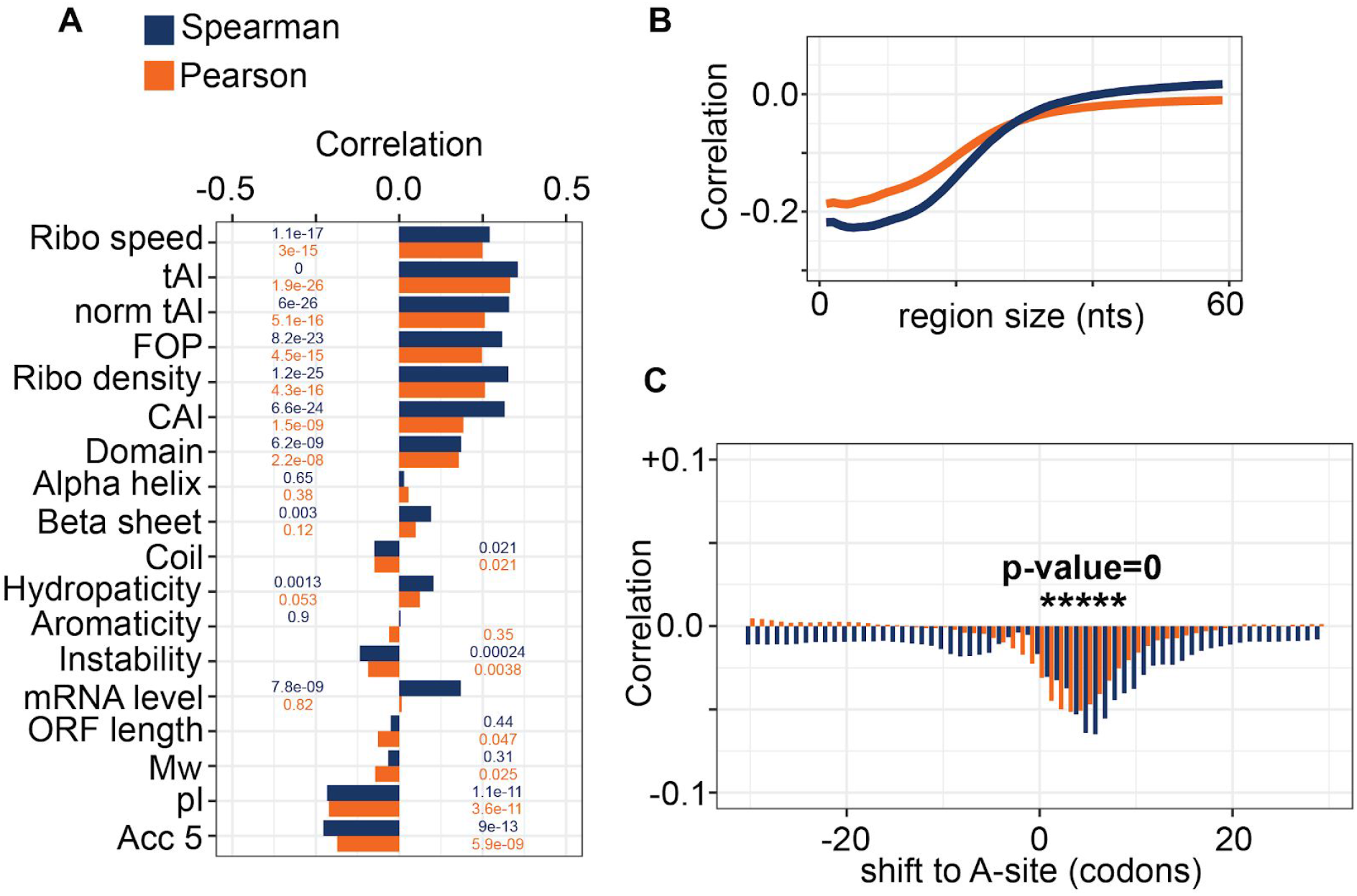
Determinants of translation elongation rate in yeast. **(A)** Correlation coefficients (p-values indicated on the bars) of SDR with features related to codon speed, biochemical properties of the encoded protein and RNA accessibility. **(B)** Correlation coefficients of the average accessibility of regions in the ORF of length indicated by the x-axis with SDR. **(C)** Correlation coefficients of SDR with the probability of regions of 20 nts starting at the position indicated on the x-axis relative to the A site to be in single-stranded conformation. Positions where the correlation coefficients are highly significant (p-value ≈ 0) are marked by asterisk. In all panels Spearman and Pearson correlation coefficients are shown in blue and orange, respectively.

Strikingly, the feature most strongly anti-correlated with SDR was the estimated isoelectric point (pI), reflecting the charge of the encoded protein. Other global features of the encoded protein such as the proportion of aromatic amino acids (Aromaticity), hydropathicity (measured by the GRAVY (GRand AVerage of hYdropathy) index (30)), molecular weight (Mw) and instability (measured by the instability index (31)) had much smaller correlation with SDR and thus with translation speed. Incorporating all of the features shown in Figure 4A into a linear model predicted better the protein synthesis rate than the ribosome density alone (correlation coefficient 0.69 vs. 0.57, Fisher’s Z test p-value = 4.94e-6). The linear model also highlighted the most explanatory features, which were (in order of the significance of their correlation coefficient being different than zero) the ribosome density (p-value < 2e-16), isoelectric point pI (p-value = 3.64e-15), codon adaptation index (p-value = 1.22e-11), molecular weight of the encoded protein (p-value = 9.83e-05), ORF length (p-value = 0.000118), GRAVY index (p-value = 0.000256), tRNA adaptation index (p-value = 0.000977), domain coverage (p-value = 0.006493) and fraction of optimal codons (p-value = 0.009106). The weights of individual features in the linear model are shown in <u>Supplementary Table 1</u>. Qualitatively similar but somewhat lower in magnitude correlation coefficients were obtained when the RNA-seq data from (4) were used in the analysis (Supplementary Figure 3).

Consistent with RPs having low translation efficiency (Figure 2D), the set of 50 genes with lowest SDR was strongly enriched in RPs (19 RP genes, hypergeometric test FDR = 4e-18 for the enrichment of the ‘Ribosome’ KEGG category). No specific biological process or cellular component was preferentially represented among the genes with highest elongation rates.

### Complex effect of RNA secondary structure on translation elongation

Although it did not significantly contribute to the linear model, the structural accessibility of translated RNAs - measured by the average probability of windows of *n* nucleotides (nts) along the ORF of being predicted in single stranded conformation - was anti-correlated with SDR for *n* up to 20-40 nts (Figure 4B). This indicates that RNAs that are highly structured are also highly translated (32). Although this seems counterintuitive, a theoretical study proposed that structural rearrangements of the mRNA during translation may serve to maintain an optimal ribosomal flux for high protein output (33). On the other hand, structural accessibility of the RNA immediately ahead of the decoded codon was significantly anti-correlated with the ribosome density on the decoded codon, as found in vitro (34, 35) (Figure 4C). Our results thus indicate that although ribosomes progress faster through unstructured regions of the ORFs, unstructured RNAs ultimately have lower translational output.

### Influence of incorporated amino acids on translation speed

A functional analysis uncovered sequence-dependent rearrangements of the nascent polypeptide in the ribosomal exit tunnel, suggesting that side chain size and charge of the incorporated amino acid impact the rate of polypeptide chain elongation, as do co-translational protein folding and interaction with chaperones (36). Indeed, our analysis provides evidence for both size and charge of amino acids affecting translation speed; negatively charged proteins are synthesized at up to ~10 fold higher rates relative to neutral and positively charged proteins whose ORFs have similar ribosome densities (Figure 5A,B). Furthermore, among non-polar amino acids, those with small side chains are associated with faster elongation, whereas the more voluminous ones have the opposite effect (Figure 5C). The amino acid charge and relative abundance of cognate tRNAs impact translation elongation rate to a similar degree.

**Figure 5.**
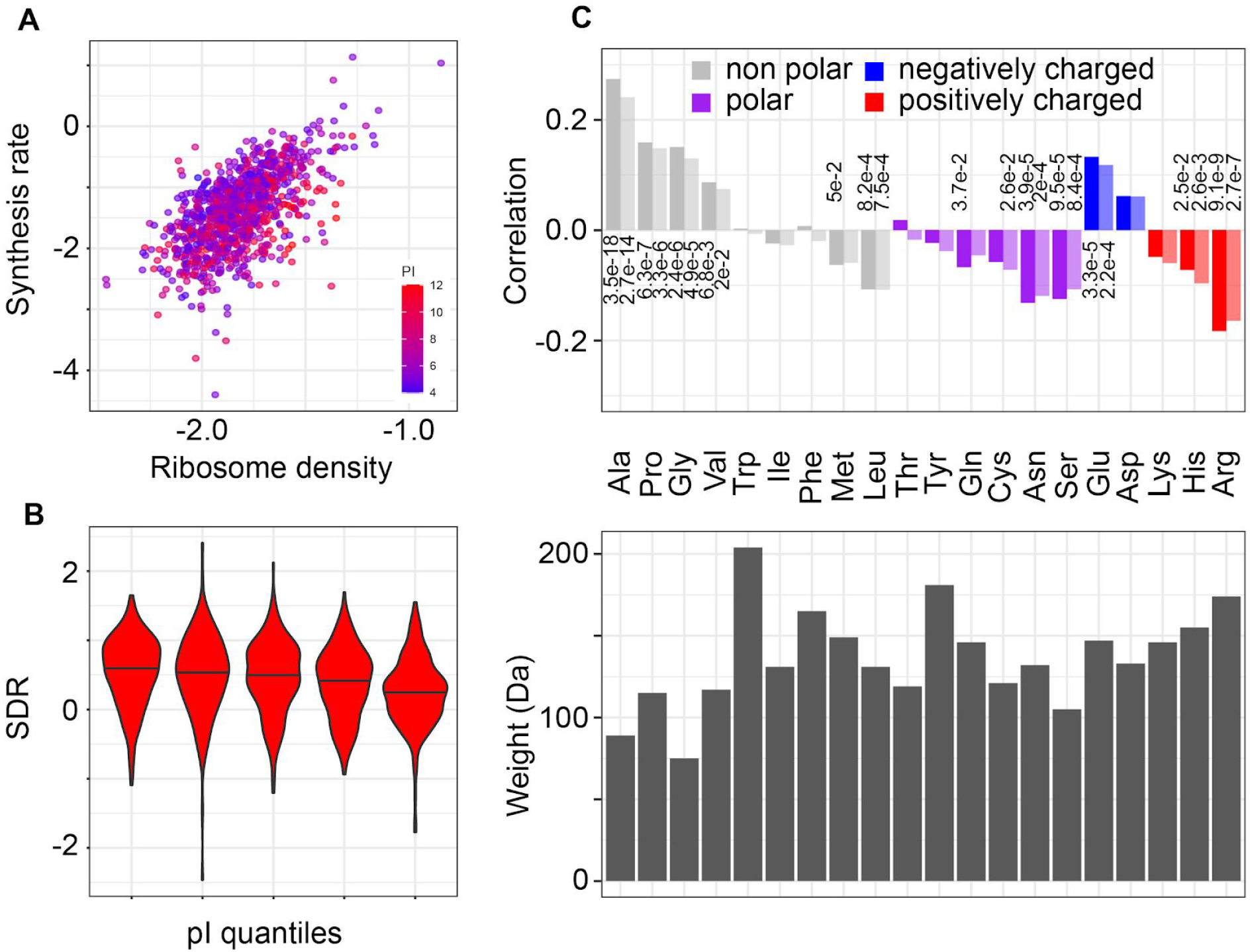
Influence of encoded amino acids on the translation elongation rate. **(A)** Positively charged proteins have low synthesis rate for the density of ribosomes on their corresponding ORFs. Each point represents an mRNA, with x and y coordinates corresponding to the ribosome density and protein synthesis rate, respectively, both on a log10 scale. The color indicates the isoelectric point of the encoded protein, red indicating proteins with high pI (positively charged) and blue proteins with low pI (negatively charged). **(B)** SDR distributions for increasing isoelectric point quantiles (left-right bins t-test p-value = 2e-8). **(C)** Pearson correlation coefficients of SDR with amino acid frequencies in the encoded proteins (top panel, only p-values < 0.05 are shown) and respective amino acids weights (bottom panel).

The explanatory power of linear models using relative frequencies of encoded amino acids or features related to tRNA abundance along with the ribosome density on the ORF were very similar (Pearson’s R 0.69 vs. 0.68). The most informative amino acids were Arg (p-value of the coefficient being different than zero in the linear fit = 2.35e-10), Pro (p-value = 1.68e-07), Ala (p-value = 1.87e-07), Glu (p-value = 1.14e-06), and Ser (p-value = 0.00861). See Supplementary Table 3 for the inferred weights of these features.

### Ribosomal proteins are translated slowly

As RP-encoding genes have a significantly higher tRNA adaptation index than other genes (Figure 6A, t-test p-value = 9.7e-49), their ORFs should be translated very efficiently. However, RPs also have a much higher isoelectric point than other proteins (Figure 6B, t-test p-value = 2.3e-47). Visualizing the tAIs of the ORFs and the pIs of proteins along with the protein synthesis rate and ribosome allocation on individual ORFs, clearly illustrates that positively-charged proteins stand out as having lower than expected protein output for the ribosome densities on the corresponding ORFs (Figure 6C, Supplementary Figure 4). This indicates that the interaction of positively-charged proteins, and in particular of RPs, with the negatively charged exit tunnel, slows down translation elongation and limits the translational output of the corresponding ORFs.

**Figure 6.**
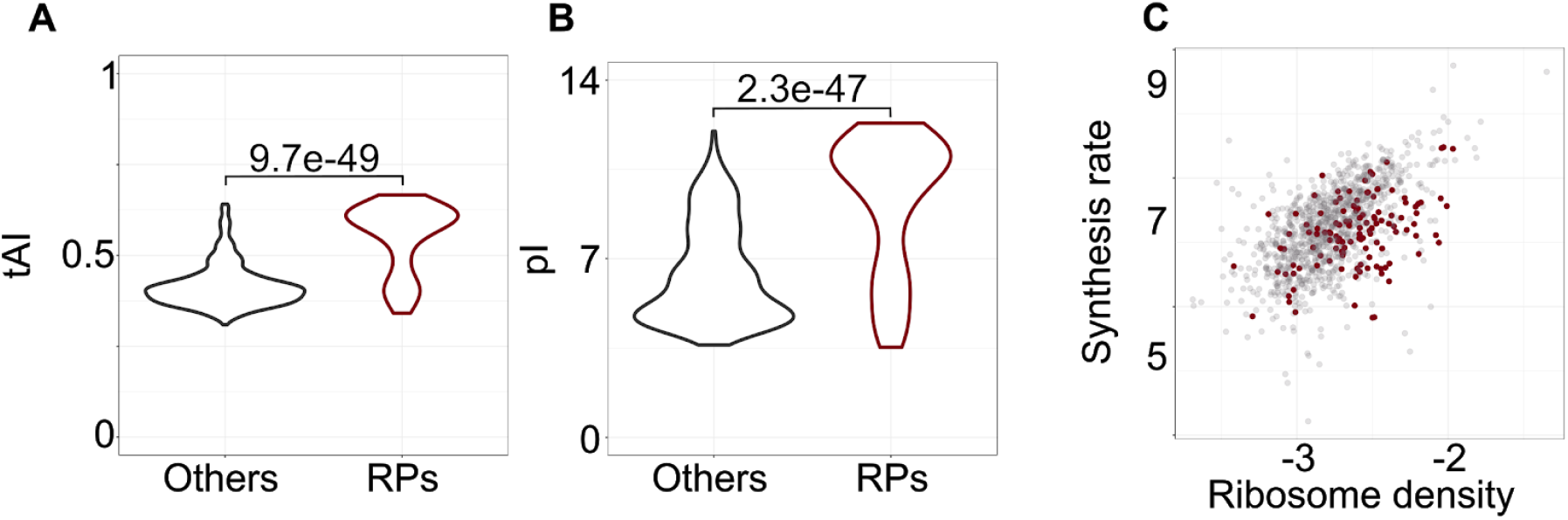
Properties of RPs (n=122) compared to all other quantified yeast proteins (n=992). Box plots of **(A)** tRNA adaptation index of individual ORFs and **(B)** isoelectric point for corresponding ribosomal and all other yeast proteins. **(C)** Protein synthesis rate (log10(peptides/s/mRNA)) as a function of ribosome density (log10(RPKM/TPM)) for transcripts encoding ribosomal proteins (brown) and all other proteins (gray).

### Perturbed translation dynamics in ribosomal protein deletion strains

Deletion of specific ribosomal protein genes has been associated with changes in translation and replicative lifespan (17, 37). In particular, deletion of *rpl7a* (*Δrpl7a* strain) led to ribosome assembly defects and overall decreased protein synthesis (measured by the incorporation of a methionine analog), whereas deletion of *rpl6a* (*Δrpl6a strain)* led to increased protein production (17). To determine how the translation parameters of individual ORFs are affected in these strains, we measured the protein synthesis rates in these strains by pSILAC and analyzed them jointly with ribosome profiling data obtained before (17). Results for individual ORFs are given in Supplementary Tables 4-6, for the wildtype control, *Δrpl6a* and *Δrpl7a* strains.

We found that accumulation of light peptides in these strains was less well explained by a linear fit compared to the wildtype strain, especially for the high-translation *Δrpl6a* strain (Supplementary Figure 5). In both mutant strains the correlation between ribosome density and protein synthesis rates was lower (Figure 7A,B) compared to the wildtype maintained in the same conditions (Supplementary Figure 6). For the *Δrpl7a* strain the decrease was due, in large part, to ORFs encoding proteins involved in starch and sucrose metabolism (FDR = 1.86e-6), glycolysis and gluconeogenesis (FDR = 0.00389), whose protein output was higher than expected for their observed ribosome densities in all of the strains in the conditions of our experiments (Figure 7B and Supplementary Figure 6). Excluding the 31 ORFs with a log10 change in ribosome density lower than −1 lead to correlation coefficients comparable with those obtained for the *Δrpl6a* strain (both Pearson and Spearman correlation coefficients = 0.41). We then compared the synthesis rate - density relationship of these strains with that of the wildtype strain analyzed in the same study (17) (Supplementary Figure 6). The ribosome density changed very little in the *Δrpl6a* strain (Figure 7C), and large, correlated changes in density and flux (more than 2 fold in either direction) were only observed for 11 ORFs. Furthermore, we did not find any ORF that was highly translated in the wildtype strain, whose protein output collapsed in the high-translation *Δrpl6a* strain, as would be expected if ribosome collisions occurred in this high-translation strain. This was not due to missing protein synthesis rate data, because the large majority (25 of 31) of ORFs with highest ribosome density and measured output in the wildtype were also measured in the *Δrpl6a* strain.

**Figure 7.**
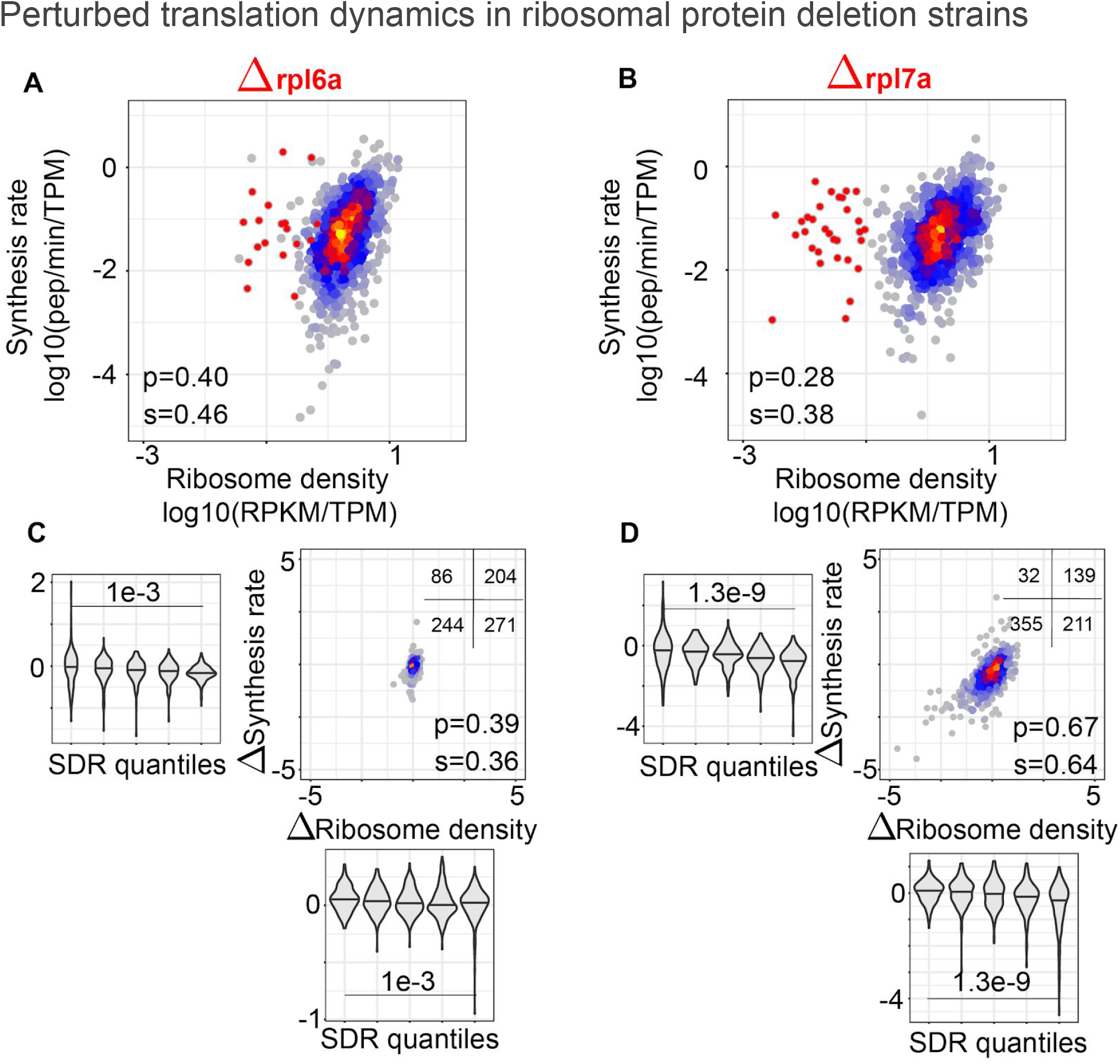
Translation parameters in yeast strains with deletions in RP genes, *Δrpl6a* and *Δrpl7a*. **(A)** and **(B)** density synthesis plots for the two strains with *Δrpl7a* strain outliers (having ribosome density < 0.01 RPKM/TPM in this strain, Supplementary Table 7) highlighted. **(C)** and **(D)** change in the ribosome density vs. change in the synthesis rate in the *Δrpl6a* and *Δrpl7a* strains compared to wildtype. The insets show the number of transcripts in each of the four quadrants of the plots. In the violin plots, the distributions of density and synthesis rate changes are shown for 5 bins of SDR values (20% of transcripts in each bin), from the lowest SDR (left-most bin) to the highest (right-most bin). P-values of the t-test comparing the mean density and synthesis rate changes between 20% transcripts with highest and lowest SDR values, respectively are shown.

In contrast, hundreds of proteins had reduced synthesis rates in the *Δrpl7a* strain relative to wildtype, with correspondingly reduced ribosome densities along ORFs (Figure 7D). The change in ribosome density was well correlated with the change in protein output (correlation coefficients = 0.64 (Spearman, p-value < 2e-16) and 0.67 (Pearson, p-value < 2e-16). The protein output of ORFs with high SDR/translation elongation rate in the wildtype strain decreased more in the *Δrpl7a* strain than the protein output of ORFs with low SDR/elongation rate. This is indeed the expected behavior in a strain with impaired ribosome biogenesis and thus with globally reduced translation initiation rates. This can be inferred from the size of the intervals between two lines of distinct translation initiation rates (dashed lines in Figure 3B) along two lines of high and low elongation rates (colored lines in Figure 3B). These results demonstrate that the analysis of protein synthesis rates and ribosome densities enables the inference of translation initiation and elongation parameters for individual genes and that these parameters can be used to uncover elements that regulate translation in individual strains and conditions.

## Discussion

Protein synthesis is a central activity in all cells, that has to be appropriately adjusted to the resources and signals that a cell experiences. The overall ribosome content of mammalian cells is strongly linked to their proliferation rate, in actively dividing cells ribosomal RNAs (rRNAs) taking up ~80% of all nucleic acids and ~15% of the biomass (38). Understanding how translation is regulated in relation to the cellular state is important, as changes in the protein synthesis capacity can lead to both cancers (39, 40) and changes in organism lifespan (17, 37, 41). Although theoretical models of biosynthetic processes have been proposed and studied for decades (10, 11, 42–45), measurements of translation dynamics across a large fraction of the transcriptome became possible only recently. Taking advantage of abundant data generated for the yeast *S. cerevisiae* and measuring protein synthesis rates with the high transcriptome coverage afforded by currently available methods, we evaluated the translation initiation and elongation rates for individual yeast ORFs.

Using additional data sets to estimate absolute protein synthesis rates as well as ribosome densities per codon, we found that the translation initiation rate varies over a ~100 fold range among yeast transcripts (Figure 3). This is consistent with an initial estimation of translation efficiency based on ribosome profiling (2) as well as with the results of a study that used these data to parametrize a whole-cell model of translation, which found that the time (5th to 95th percentile) between initiation events on individual mRNAs is from 4 to 293 seconds (44). In contrast, a narrower range of variation, ~11 fold (1st to 99th percentile), was inferred in the initial analysis of the ribosome profiling data that we also used here (4), as well as in a subsequent study of a more limited set of proteins (14). It was suggested that inaccuracies in estimation of mRNA expression levels could account for discrepancies in estimates of translation efficiency from ribosome profiling. However, we found that a much wider range of variation in translation initiation rates, of ~150 fold (Supplementary Figure 7), was necessary to explain the protein synthesis rate data, even when the mRNA levels used on the analysis were obtained with a protocol designed to minimize these inaccuracies (4). The similar results obtained based on mRNA level estimates with two sequencing protocols is perhaps not surprising, as the 3’ end bias of the data was also similar (Supplementary Figure 2) and the transcript abundance estimates showed limited systematic differences between the two data sets (Supplementary Figure 8). It is unlikely that the wider range in translation initiation rate is due to error in estimating the protein synthesis rates because our analysis only included ORFs for which peptide accumulation was well described by a constant accumulation rate. The selection of transcripts for analysis in different studies may account for some of the reported differences in the range of rate variation, as for e.g. the study of (14) used only ORFs of at least 200 codons and with a minimum ribosome coverage of 10 per site. This amounted to 894 ORFs, of which 826 are also covered by our analysis. However, our analysis includes 290 additional ORFs, some with relatively low translation. In spite of this, the mean initiation and elongation rates in our data are quite close to those reported in the (14) study, namely mean waiting time between initiation events of ~25*s* compared to the 8s median, and elongation rates of 2.63 amino acids/*s* compared to 5.6 amino acids/*s*. More importantly, previous studies did not measure protein outputs directly, but rather estimated initiation and elongation rates from ribosome densities. This can be done up to a constant scale factor, which was assumed to be identical between genes and set such as to achieve a specific target elongation rate towards the 3’ end of the ORF (14).

Our data indicates, however, that there are substantial differences in protein output of ORFs with similar ribosome densities, underscoring the importance of direct measurements of protein synthesis rates to analyze the dynamics of translation.

The translation parameters of short ORFs, many of which encode ribosomal proteins, have been the topic of much discussion (4, 44). The high ribosome density observed on short ORFs (19) has been attributed to their being evolutionarily optimized for protein output through high rate of translation initiation (44). As the high codon adaptation index exhibited by these ORFs would predict fast elongation and thereby low ribosome density (44), high ribosome density on short ORFs has also been interpreted as evidence for initiation being the main determinant of ribosome density. Consistently, we also found a small but significant correlation between ORF length and the principal component of the protein synthesis rate - ribosome density scatter, which is indicative of the translation initiation rate (Figure 3 and Supplementary Figure 9). However, our results reveal a more complex picture, showing that the charge of the encoded protein is an important determinant of ribosome flow, ORFs with similar overall ribosome density differing by up to ~20 fold in protein output. This effect is not captured by models that assume that the rate of elongation depends only on the tRNA availability-dependent decoding speed at the A site of the ribosome. Indeed, we demonstrated that protein synthesis rates can be predicted with significantly higher accuracy when taking into account global features of the encoded protein such as the pI than when using solely the ribosome density.

Overall, we found that the rate of elongation varies ~20 fold among yeast ORFs, less than the rate of initiation. As the determinants of translation elongation rate are actively debated (9, 29, 44, 46, 47), we evaluated their relative contributions in our data. We further included in our study yeast strains with globally perturbed translation through ribosomal protein gene deletions. We found no evidence that translation elongation severely curbs protein output, either in the exponentially growing *BY4741* yeast strain, or in the *Δrpl6a* and *Δrpl7a* deletion strains, the first with higher and the second with lower overall protein synthesis rate compared to the *BY4741* wildtype. Rather, several lines of evidence point to evolutionary optimization of ORF sequences to maintain appropriate ribosome flux and minimize the chance of ribosome collision. For instance, ORFs with high protein output have high rates of translation initiation and at the same time a high codon adaptation index. This is predicted to enable fast elongation, as optimal codons will be rapidly found by cognate tRNAs that are in highest abundance. Our data provides transcriptome-wide evidence for the high elongation rates of highly expressed ORFs. Thus, although initiation rates vary over a wide range, the protein output increases in parallel with the ribosome density, without the latter reaching saturation. Moreover, the density of RNA secondary structure predicted in the ORF was positively correlated with the translation elongation rate, not negatively correlated, as would be expected if RNA structure were to hinder translation. This indicates that the RNA structure may also help maintaining the flux of ribosomes along the ORF to minimize ribosome collisions, as proposed in a previous study (33). Our results do not exclude ‘controlled’ ribosome stalling at specific positions, such as on upstream open reading frames (48), or at codons for which cognate tRNAs are limiting in specific conditions, where active regulatory mechanisms are used to modulate the output of specific ORFs (47). They also do not exclude clearance of the ribosomes from the 5’ end of transcripts effectively reducing the initiation rate to some extent (the concept of 5’ ramp (14)). Rather, our data supports the notion that ORF have undergone evolutionary selection to minimize the chance of ribosome stalling due to imbalanced initiation and elongation rates.

That the charge of the translated protein affects the rate of translation elongation has been observed before (7, 8, 14, 49), and has been attributed to variation in the “friction” of the polymeric chain with the ribosomal exit tunnel. This effect is most marked for the positively charged RPs, whose elongation rate is low relative to other proteins whose transcripts have similar ribosome densities. Our results thus provide a rationale for the previous observation that the ratio of protein to mRNA molecules is markedly lower for RP-encoding compared to other genes (50). The interaction of RPs with the negatively charged rRNAs likely imposes a strong selection pressure for positive charge on RP genes (51), which in turn sets an upper bound on the rate of translocation of the polypeptide chain through the ribosome channel. It will be interesting to explore whether this slower elongation rate may have as side effect an increased translation fidelity of these very abundant proteins (52).

Over ten years ago it has been discovered that protein folding takes place already co-translationally and that helices can fold within the ribosome exit tunnel (53). A recent study further suggested that non-optimal codons drive effective co-translational folding of α-helices and β-sheets (26). Although our results are consistent with these conclusions, they indicate that the positive correlation of translation speed with high density of protein domains is not limited to particular secondary structure elements.

All of the distinct features that we analyzed here, namely tRNA/codon usage, structure accessibility of the RNA and protein charge, have small and comparable correlation with elongation rate. Altogether, they explain ~½ of the variance in elongation rate. This indicates that more detailed models that also include positional features (14) as well as more accurate ribosome coverage profiles (54) will be necessary to improve the prediction of translation dynamics. Our measurements of protein synthesis rates in multiple yeast strains with different translation capacity provide an ideal test bed for new models.

## Materials and Methods

### Simulations

All simulations of the TASEP model have been performed with C++ code developed inhouse and available upon request. The size of the ribosome footprint has been set to 10 codons.

### Analysis of poly-A selected RNA-seq

Reads from the fastq files associated with the publication of (17) have been trimmed with cutadapt (55) with parameters ‘—error-rate 0.1—minimum-length 15—overlap 1’, first of the 3’ adapter (TGGATTCTCGGGTGCCAAGG) and then of the poly-A tail (‘adapter’ = (A)_50_). Resulting sequences (of at least 15 nts) were then mapped to yeast ORFs obtained from the yeastgenome.org (56) database (URL https://downloads.yeastgenome.org/sequence/S288C_reference/orf_dna/), with the bowtie2 (57) aligner, version 2.2.9 with parameter ‘-q’ (fastq format). For each read, the best alignment reported by bowtie2 was used, and expression levels for each ORF, expressed in reads per kilobase per million (RPKM) were calculated by dividing the number of reads mapped to the ORF by library size and ORF length, then multiplying by 10^6^. For each ORF, the expression level used in the analysis was the average computed from three replicates.

### Analysis of ribosome profiling data

Ribo-seq data have been downloaded from the GEO database (58) (accession: GSE53313). Reads from raw fastq files were trimmed with cutadapt (3’ adapter: TCGTATGCCGTCTTCTGCTTG) with parameters ‘—error-rate 0.1—minimum-length 15—overlap 1’. The first 8 nts corresponding to random barcodes were then trimmed as well and the remainder of the sequence was first aligned to ribosomal RNAs with bowtie2 version 2.2.9, parameters ‘-q’ and ‘—un’ to indicate the fastq format of the input file and to obtain also the unmapped reads. The latter were then aligned to a database consisting of all yeast ORFs extended by 200 nts upstream and downstream, to be able to reconstruct full-length ribosome profiles along the ORFs. To allow for the possibility of closely spaced ORFs which would lead to reads mapping in an overlapping manner to the two genes, we extracted up to two best mappings, with bowtie2 parameter ‘-k 2’. The positions of the ‘A’ site of ribosomes in individual reads were inferred as in (4).

### Analysis of protein synthesis rates

For the pSILAC experiment, the *S. cerevisiae* strain *BY4741* was grown as described (22). Briefly, synthetic medium containing 2% glucose, yeast nitrogen base (6.7 gm/l), and dropout medium (2 g/l) containing all the amino acids except L-lysine was prepared. Initially, a pre-culture of yeast was grown at 30 °C, 200 rpm, in three biological replicates obtained by inoculating three different colonies in 5 ml of heavy SILAC synthetic medium containing heavy L-Lysine-2HCl, 13C6, 15N2 (ThermoFisher# 88209) at a final concentration of 30 mg/l. The pre-culture step was repeated one more time so that all the proteins became tagged with heavy isotope. The pre-culture thus obtained was used to grow cells at optical density of A600 = 0.4 in 200 ml. At this point, cells were centrifuged, washed twice with light SILAC synthetic medium containing light L-Lysine-2HCl (ThermoFisher# 89987) at concentration of 30 mg/l and transferred to 200 ml of light SILAC media. Cells were harvested at 0, 5, 15, 30, 60, 120, 180 minutes for the mass spectrometric analysis.

Cells were lysed in a buffer containing 1% sodium deoxycholate, 0.1M ammonium bicarbonate and 10mM TCEP using strong ultra-sonication (Bioruptor, 10 cycles, 30 seconds on/off, Diagenode, Belgium). Samples were heated to 95°C for 10 minutes and after cooling, the protein concentration was determined by the BCA assay (Thermo Fisher Scientific), using a small sample aliquot. 50 μg of protein were alkylated with 15 mM chloroacetamide for 30 min at 37 °C and incubated with sequencing-grade modified trypsin (1/50, w/w; Promega, Madison, Wisconsin) overnight at 37°C. After acidification using 5% TFA, precipitated detergent was removed by centrifugation (14k rpm, 5 min). Peptides were desalted on C18 reversed-phase spin columns according to the manufacturer’s instructions (Microspin, Harvard Apparatus) and dried under vacuum.

The setup of the μRPLC-MS system was as described previously (59). Chromatographic separation of peptides was carried out using an EASY nano-LC 1000 system (Thermo Fisher Scientific), equipped with a heated RP-HPLC column (75 μm x 37 cm) packed in-house with 1.9 μm C18 resin (Reprosil-AQ Pur, Dr. Maisch). Aliquots of 1 μg total peptides were analyzed per LC-MS/MS run using a linear gradient ranging from 95% solvent A (0.15% formic acid, 2% acetonitrile) and 5% solvent B (98% acetonitrile, 2% water, 0.15% formic acid) to 30% solvent B over 90 minutes at a flow rate of 200 nl/min. Mass spectrometry analysis was performed on Q-Exactive HF mass spectrometer equipped with a nanoelectrospray ion source (both Thermo Fisher Scientific). Each MS1 scan was followed by high-collision-dissociation (HCD) of the 10 most abundant precursor ions with dynamic exclusion for 20 seconds. Total cycle time was approximately 1 s. For MS1, 3×10^6^ ions were accumulated in the Orbitrap cell over a maximum time of 100 ms and scanned at a resolution of 120,000 FWHM (at 200 m/z). MS2 scans were acquired at a target setting of 10^5^ ions, accumulation time of 50 ms and a resolution of 15,000 FWHM (at 200 m/z). Singly charged ions and ions with unassigned charge state were excluded from triggering MS2 events. The normalized collision energy was set to 27%, the mass isolation window was set to 1.4 m/z and one microscan was acquired for each spectrum.

The acquired raw files were imported into the Progenesis QI software (v2.0, Nonlinear Dynamics Limited), which was used to extract peptide precursor ion intensities across all samples applying the default parameters. The generated mgf files were searched using MASCOT using the following search criteria: full tryptic specificity was required (cleavage after lysine or arginine residues, unless followed by proline); 3 missed cleavages were allowed; carbamidomethylation (C) was set as fixed modification; oxidation (M) and heavy SILAC (K8) were applied as variable modifications; mass tolerance of 10 ppm (precursor) and 0.02 Da (fragments). The database search results were filtered using the ion score to set the false discovery rate (FDR) to 1% on the peptide and protein level, respectively, based on the number of reverse protein sequence hits in the datasets. The relative quantitative data obtained were normalized and statistically analyzed using our in-house script (SafeQuant) as above (59).

The protein synthesis rates have been obtained from the slope of linear regression constrained to 0 at time point 0. This analysis was performed with the *lm()* function of R (version 3.4.2). For each regression, R-squared values have been recorded for further analysis (Fig. 2C).

### Scaling mRNA copy numbers, protein synthesis rates and ribosome densities

To obtain absolute protein synthesis rates, the relative rates obtained from pSILAC were scaled, using the known values of the number of protein molecules per yeast cell (21), the doubling time of yeast, and the average half life of yeast proteins (22). Furthermore, as the proteomics experiment does not capture all proteins, the uncaptured fraction had to be taken into account. The fraction of captured proteins has been approximated as follows: reads from ribosome footprints were used to compute normalized ORF abundances in the Ribo-seq data (RPKMs). The total abundance of translated ORFs that were not captured in the proteomics data, relative to all of the ORFs captured in Ribo-seq was used as the fraction of uncaptured protein. The steady-state level of protein per cell should be given by the ratio of synthesis and degradation rates. The synthesis rate can thus be calculated as the product of the steady-state level of protein per cell and the degradation rate. The latter is the result of two processes, protein degradation and dilution due to cell growth. This leads to the following expression for the average synthesis rate of the captured fraction

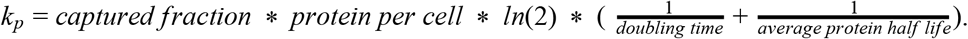

To obtain synthesis rates for individual ORFs, we multiplied the total synthesis rate of the captured fraction by the relative synthesis rate inferred by fitting the light peptide accumulation in pSILAC

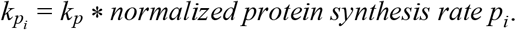

To infer absolute densities of ribosomes per codon, we determined the first principal component of the absolute ribosome densities from (19) relative to our estimates based on RPFs (Fig. 2A). We used this first principal component to map relative densities we obtained based on RPF and mRNA-seq reads to ribosomes densities per codon for each ORF.

Absolute abundances of mRNA molecules per cell were obtained by rescaling the relative numbers inferred from RNA-seq to obtained a total of 40,000 transcripts per cell, as found in previous work (23, 24).

### Computation of mRNA/protein features

Protein features used in the analysis of translation elongation rate have been downloaded from the yeast genome database at the following link https://downloads.yeastgenome.org/curation/calculated_protein_info/protein_properties.tab. The tRNA adaptation index (tAI) and normalized tAI (ntAI) have been computed as in (25) and (26) with a custom python script. The RNAplfold tool from ViennaRNA package (60) (version 2.1.8) was used with default parameters to estimate structural accessibility along ORFs. For each ORF, the average accessibility of windows of a specified size has been calculated.

We analyzed the following features:

- Molecular weight (Mw) of the protein in daltons (Da).
- Isoelectric Point (pI): the pH at which the protein does not carry net electric charge.
- *GRand AVerage of hYdropathicity* (GRAVY Score): the average of hydropathy values of all amino acids in the protein (30).
- *Aromaticity Score*: the frequency of aromatic amino acids, Phe, Tyr and Trp.
- *Codon Adaptation Index* (CAI): measures the bias of codon usage in a coding sequence in respect to a reference set of genes (28). It is defined as the geometric mean of the weights over all codons in the sequence (*L*):

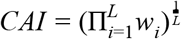

where the weight of each of codon is computed from the reference sequence set, as the ratio between the observed frequency of the codon *f_i_* and the frequency of the most frequent synonymous codon *f_i_* for that amino acid.

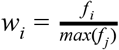

with *i, j* synonymous codons.
- *tRNA Adaptation Index* (tAI): measures the adaptation of each transcript to the pool of tRNAs (25). Similarly to CAI, tAI is the geometric mean of weights associated to each codon

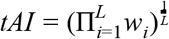

where

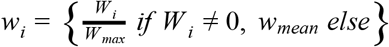

and

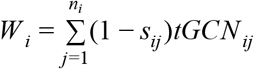

with *n_i_* number of tRNA isoacceptors, *tGCN_ij_* gene copy number of tRNA *j*-th recognizing *i*-th codon and *s_ij_* is a selective constraint of codon-anticodon coupling.
- *normalized tRNA Adaptation Index* (ntAI): normalized version of tAI based on the codon usage in the transcriptome (26), it has a similar form to tAI, but with *W_i_* being scaled by the normalized codon expression in the following way: *U_i_* is the usage of codon *i* taking into account the abundance of individual transcripts

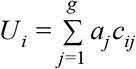

with *a_j_* being the transcript abundance of gene *j* and *c_ij_* is the number of occurrences of codon *i* within the ORF of the gene *j*. The usage of codon *i* is then defined as

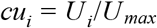

and finally the weights that used in the calculation of the tAI are defined as

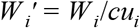

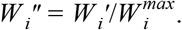 The factor 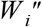substitutes *W_i_* in the formula for tAI.
- *Fraction of OPtimal codons* (FOP): fraction of optimal codons in the ORF (27). The optimal codon for an amino acid is the codon most used to encode the amino acid in the ORFs encoding the top expressed proteins.
- *Instability Index*: measure of protein half-life estimated based on the dipeptide composition of the protein (31).
- *Domain coverage*: fraction of protein covered by Pfam domains predicted by InterPROScan (61)
- *alpha-helix, beta-sheet, coil*: fraction of the protein sequence involved in the indicated types of secondary structures predicted by PSIPRED (62).
- *Accessibility 5 nts*: average probability of finding a window of size 5 nucleotides in an open conformation predicted with RNAplfold from ViennaRNA package (60).

### Enrichment tests

Gene Ontology and KEGG pathway enrichments have been performed through the STRING database (63).

## Author contributions

AR and MZ designed the research. NDN and AR performed the analysis of RPF and RNA-seq data. NM, EA and AS performed the proteomic experiments. EA and AR analyzed the proteomic data. AR, AS and MZ wrote the manuscript.

## Acknowledgements

AR would like to thank Joao C. Guimaraes for discussions and knowledge sharing that helped in developing the project.

## Supplementary Figures

**Supplementary Figure 1.**
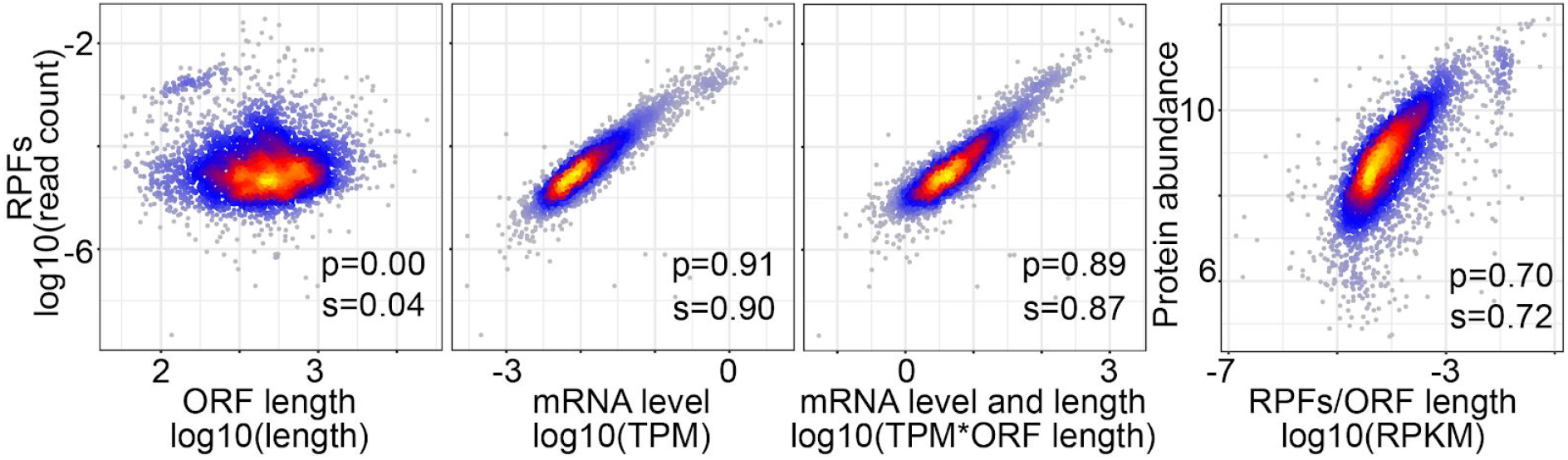
Relationship between the number of RPFs, the mRNA levels and ORF lengths inferred from the mRNA-seq and Ribo-seq data from (4). Counterintuitively, the mRNA level alone explains more of the variance in RPF numbers than the combination of mRNA level and ORF length.

**Supplementary Figure 2.**
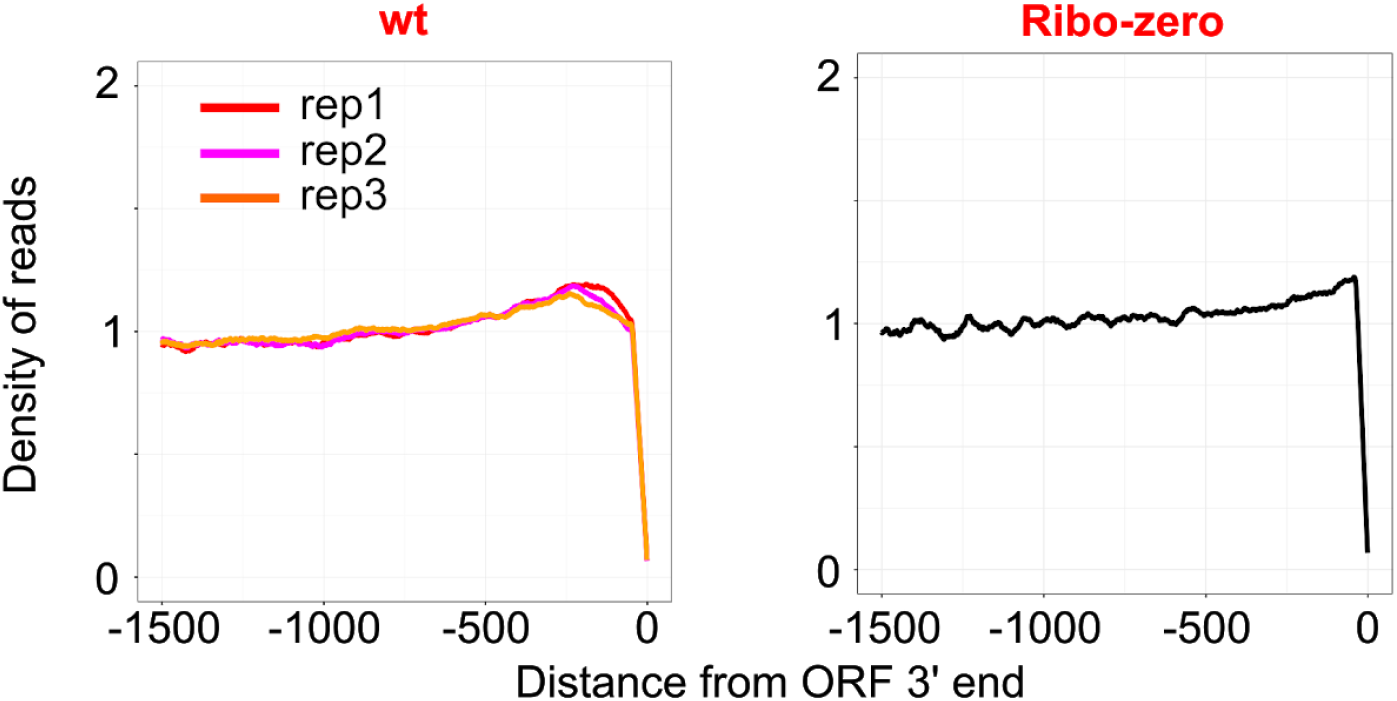
Relative coverage of transcripts by reads obtained from sequencing oligo-dT selected RNAs, as a function of the distance from the stop codon (ORF end). The coverage of each position in individual ORFs by reads was normalized by the average coverage over the entire ORF, then ORFs were anchored at the 3 end, and, for each position relative to ORF end, the mean relative coverage was computed, based on all ORFs that extended at least that far upstream from the stop. Values are shown relative to the mean coverage per position along the ORF.

**Supplementary Figure 3.**
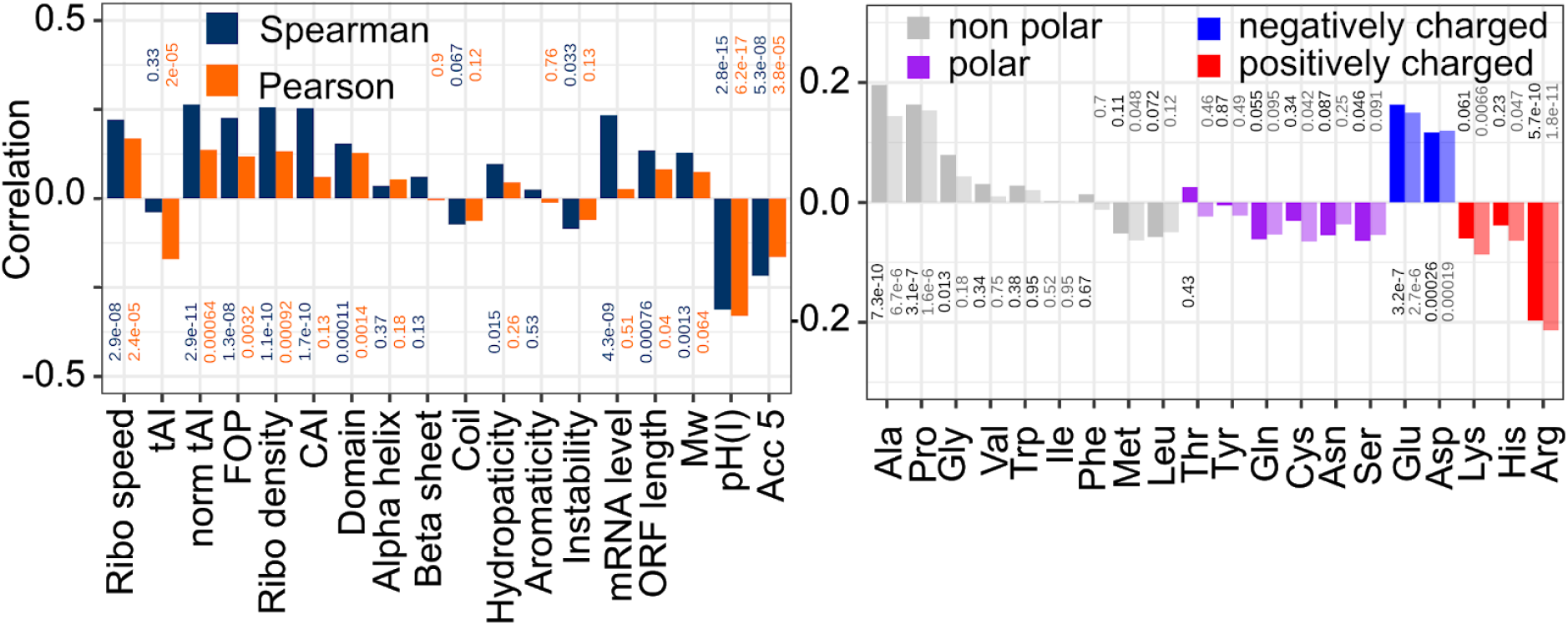
Correlation coefficients of SDR with features related to codon speed, biochemical properties of the encoded protein and RNA accessibility (left) or amino acid frequencies in the encoded proteins (right), based on Ribo-seq and RNA-seq data from (4). Spearman (blue in left and dark shade in right panels) and Pearson (orange in left and light shade in right panels) correlation coefficients are indicated on the y-axis, corresponding p-values are written on the plots. Ribo-seq and Ribo-zero data sets from (4) were used to obtain the ribosome densities, the pSILAC data generated in this study was used to compute protein synthesis rates.

**Supplementary Figure 4.**
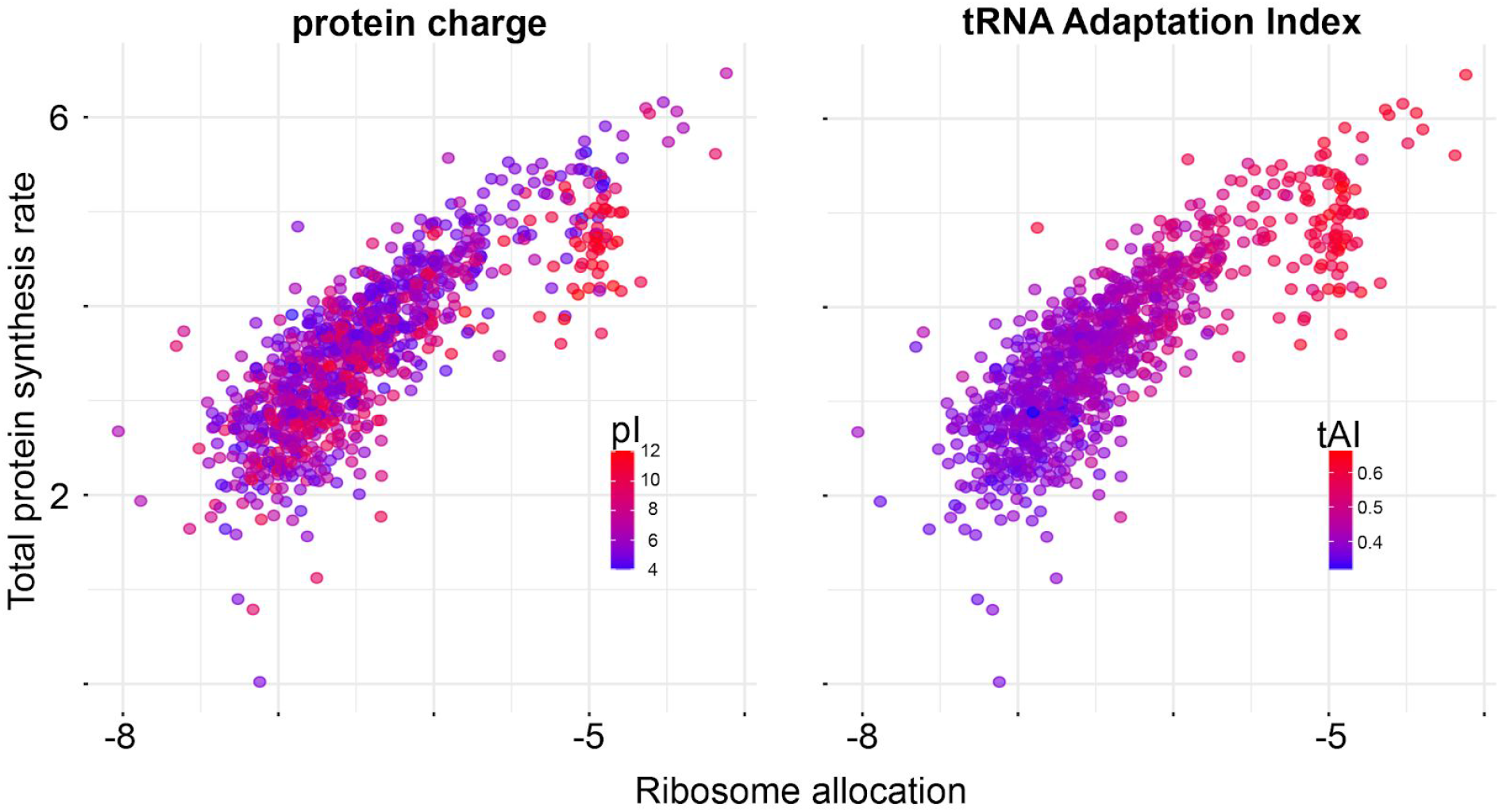
Total protein synthesis rate in function of the ribosomes allocated to each mRNA species. Each point represents a transcript, the color indicating isoelectric point of the encoding protein (left) or the tRNA adaptation index (right).

**Supplementary Figure 5.**
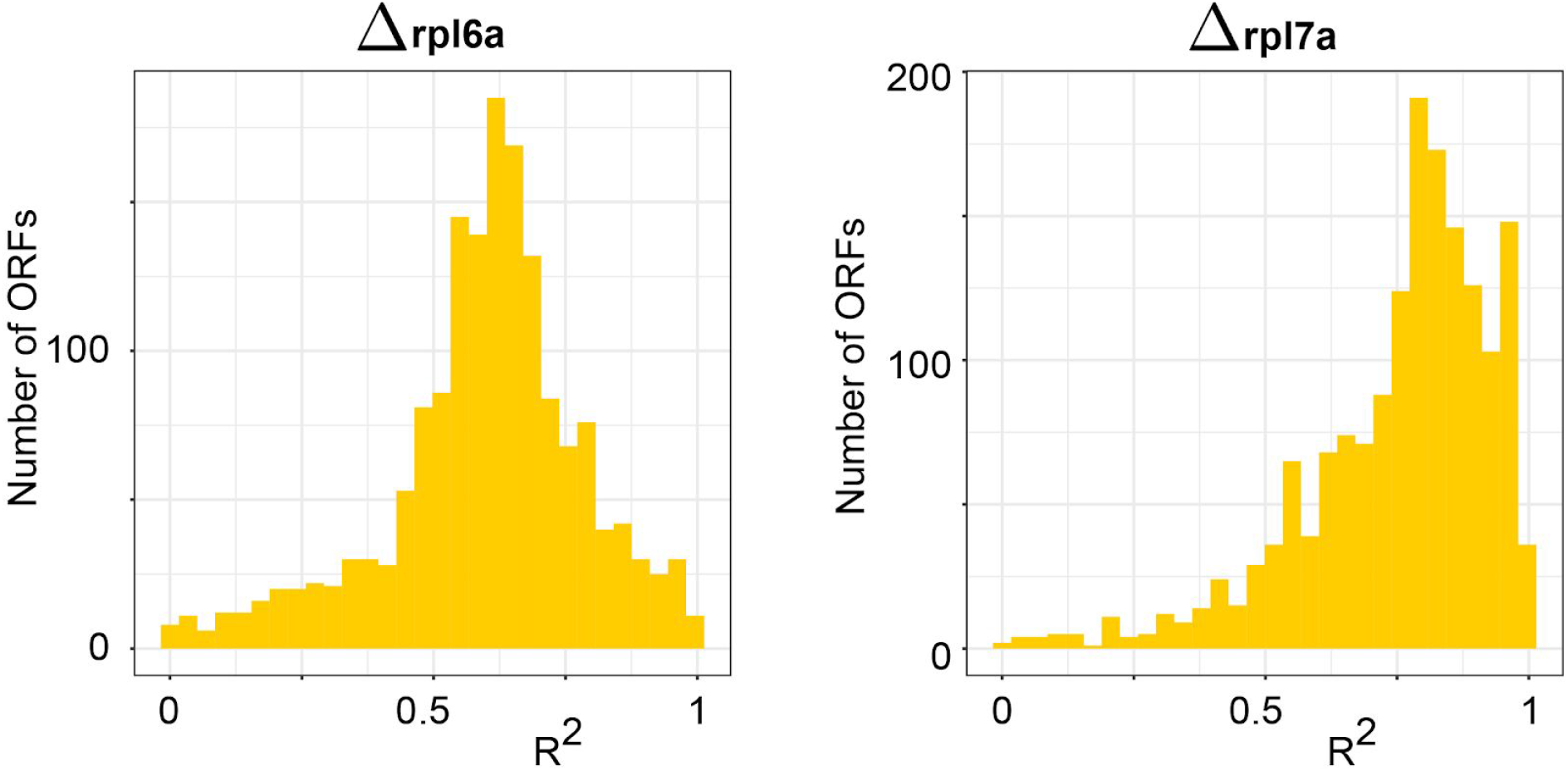
R^2^ values for the estimated protein synthesis rates in *Δrpl6a* and *Δrpl7a* strains.

**Supplementary Figure 6.**
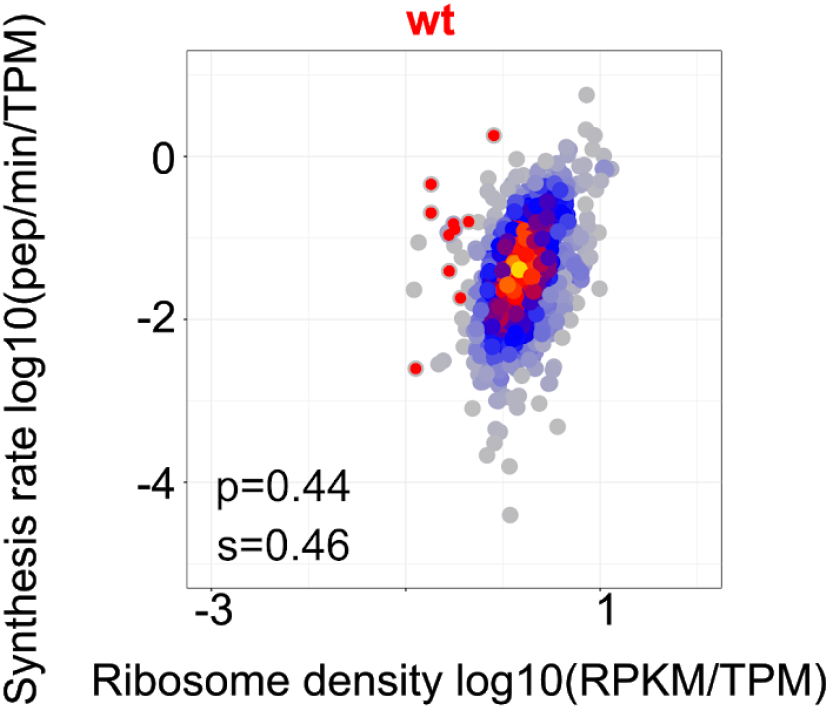
Density-synthesis plot for wt with outliers, whose ribosome densities were lower than 0.01 RPKM/TMP in the *Δrpl7a* strain, highlighted in red (see also Figure 7).

**Supplementary Figure 7.**
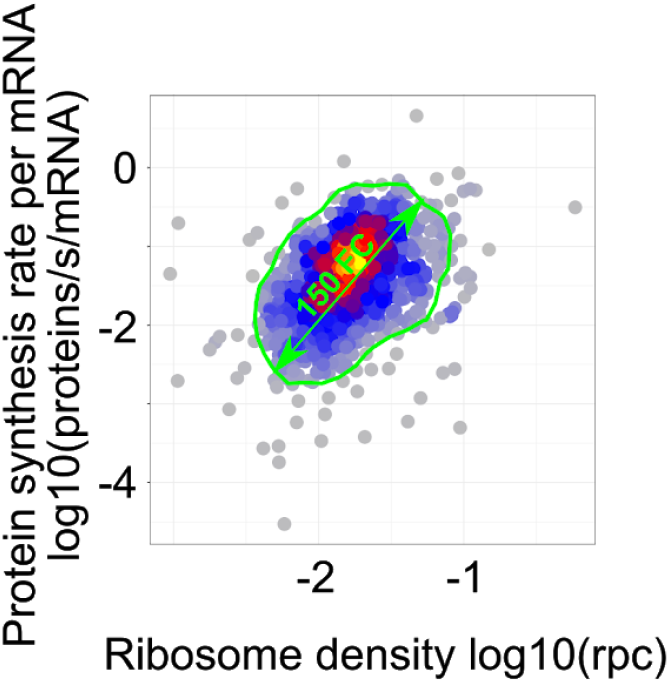
Absolute density-synthesis plot based on ribosome footprints and RNA-seq (Ribo-zero) data from (4) (see also Figure 3).

**Supplementary Figure 8.**
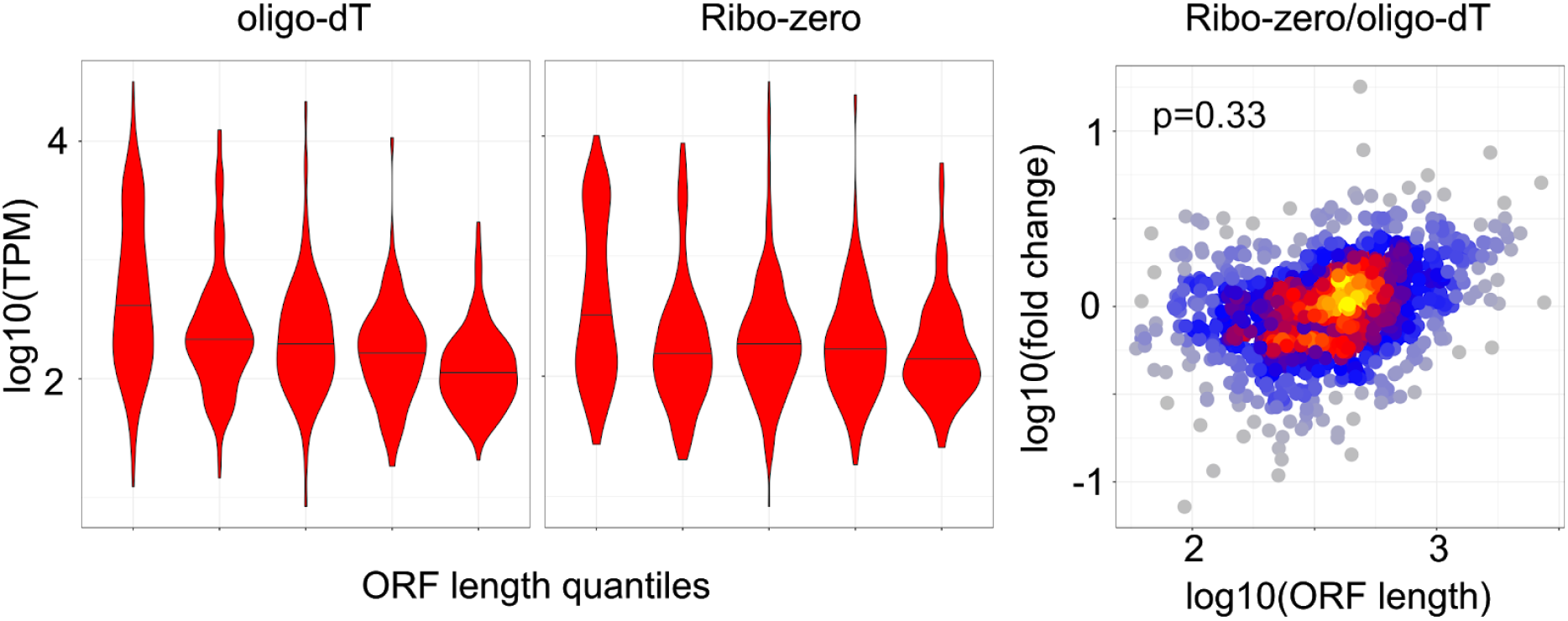
ORF-length-dependent bias in transcript capture rate with Ribo-zero vs. oligo-dT selection. The plots show the distribution of inferred expression levels of ORFs in bins of ORF length, each bin containing 20% of the transcripts. Bins are sorted from smallest (left) to largest (right). The log10 ratio of expression levels estimated with Ribo-zero relative to oligo(dT), as a function of ORF length. The bias is apparent for long ORFs, whose expression estimates are systematically higher with Ribo-seq than with oligo(dT).

**Supplementary Figure 9.**
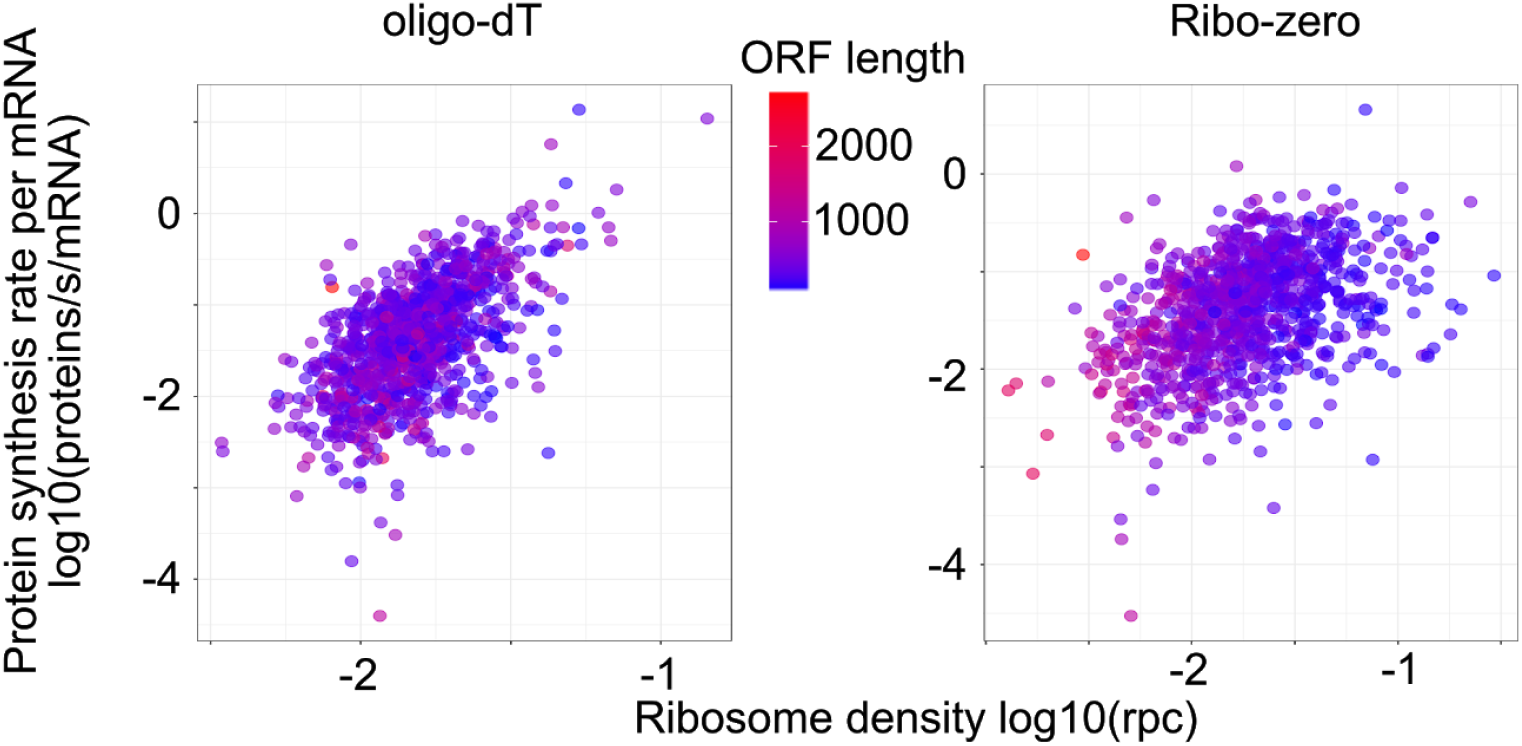
Influence of ORF length on the protein synthesis rate - ribosome density relationship. The length of corresponding ORFs (in nucleotides) is indicated on the color scale. ORF length is positively correlated with the coordinate of the ORF along the first principal component of the synthesis - density relation. Pearson correlation coefficients are −0.09 (p-value = 0.0059) in the oligo-dT-based analysis and −0.32 (p-value < 2e-16) in the analysis based on the Ribo-zero protocol.

## Supplementary Tables

**Supplementary Table 1.** Ribosome densities and synthesis rates with RPF from (4) together with oligo-dT RNA-seq from (17) for wild type condition.

**Supplementary Table 2.** Ribosome densities and synthesis rates with RPF and Ribo-zero from (4) for wild type condition.

**Supplementary Table 3.** Estimated features from the linear models.

**Supplementary Table 4.** Ribosome densities (based on data from (17)) and synthesis rates for wild type condition.

**Supplementary Table 5.** Ribosome densities (based on data from (17)) and synthesis rates for *Δrpl6a* mutants.

**Supplementary Table 6.** Ribosome densities (based on data from (17)) and synthesis rates for *Δrpl7a* mutants.

**Supplementary Table 7.** Outliers, with log10 decrease in ribosome density < −2 in *Δrpl7a* mutants.

